# A novel HER2 protein identification methodology in breast cancer cells using Raman spectroscopy and Raman imaging: an analytical validation study

**DOI:** 10.1101/2024.07.04.602049

**Authors:** H. Abramczyk, J. Surmacki, M. Kopeć

## Abstract

**Background:** Conventional assays such as immunohistochemistry (IHC) and *in situ* hybridization (ISH) used in clinical procedures for quantification of the human epidermal growth factor receptor-2 (HER2) status in breast cancer have many limitations. Our results suggest that a new Raman method may improve specificity that will result in better patients selection for HER2 targeted therapy. In the current study, we have used HER2 expression in a broad range of breast cancer phenotypes to explore the potential utility of a novel immunodetection technique, using Raman spectroscopy and Raman imaging combined with artificial intelligence models.

**Methods:** The expression of HER2 protein in different cancer subtypes was evaluated using Raman methodology to test correlations with currently used quantitative protein analysis for a broad range of cancer subtypes based on the expression levels of estrogen and progesterone receptors (ER and PR), HER2, cytokeratins and claudins. In the current study, five breast cancer cell lines: MCF-10A, MCF-7, MDA-MB-231, HTB-30 (SK-BR-3) and AU-565 were tested.

**Results:** The correlations between Raman method and conventional HER2 testing methodologies (IHC and ISH) have been tested. Raman measurements showed a strong linear correlation (*p*= 0.05, R^2^=0,9816) with IHC analysis in the studied breast cell lines: MCF-10A, MCF-7, MDA-MB- 231, HTB-30 (SK-BR-3) and AU-565 representing normal, non-tumorigenic epithelial cells, triple-positive breast carcinoma and triple-negative breast cancer cell lines.

**Conclusions:** Analytic testing of Raman spectroscopy and Raman imaging demonstrated that this method may offer advantages over currently used diagnostic methodologies. The broad range of HER2 expression on the surface of human breast normal and cancer cells from triple negative to triple positive cell lines were studied. Our data demonstrate that Raman based methods for HER2 quantitation of HER2 may offer significant progress in patient selection for HER2 targeted therapy over conventional HER2 identification. Further studies of HER2 determination by Raman methods with *in vivo* cells and *ex vivo* tissue are warranted.

## Introduction

When a virus attacks our body, the entire immune system is alerted through chemical signals of a generated protein. When cancer invades our body the warning alert is less obvious because cancer cells don’t trigger such fierce immune responses as for viruses. The mutated cancer cells are still similar to healthy cells and the immune system doesn’t recognize the distinction between them allowing the cancer cells continuing to grow, divide and spread throughout the body.

To detect subtle genomic and posttranslational modifications of proteins, DNA, RNA, lipids in cells and tissues and cellular environment that occur due to the development of a cancer we must use proper tools to measure biochemical alterations that are specific for cancer. The characteristic biochemical profiles are called cancer biomarkers that are measured as an indicator of the risk of cancer, occurrence, or patient outcome. ^1^ ^2^ ^3^ ^4^ ^5^

An example of a cancer biomarker is the HER2 gene that makes HER2 protein. Normal tissues have a low amount of HER2 protein that helps control cell division and growth. Aberrant extra copies of this gene (amplification) may lead to an excess (overexpression) of HER2 protein, which causes cells to grow more quickly. Overexpressed HER2 is reported in 20% of breast cancer and in some ovarian and gastric cancers. ^6,7^ ^8^ ^9^ ^10^ ^11^ ^12^ ^13^ ^14^ ^15^ ^16^ ^17^ ^18^

The HER2 protein (also called HER2/neu or ErbB2) is a cellular receptor that is responsible for translating signals from outside the cell into signals within the cell. The routine oncological test for all breast cancers is determination of HER2 status.

HER2 belongs to a group of receptor tyrosine kinases (RTKs) and consists of an extracellular domain that includes four subdomains (I-IV) ^19^, a single helix transmembrane lipophilic segment, and an intracellular region that contains a tyrosine kinase domain (TKD) and a carboxyl (C-) terminal tail (Fig.1) ^20^. RTKs can be activated by ligand-dependent and ligand-independent mechanisms. No ligands for HER2 have yet been identified yet. ^21^ ^22^ and dimerization with any of the other three subdomains is considered to activate HER2. ^23^ The dimerization in the extracellular region of HER2 induces intracellular conformational changes that trigger tyrosine kinase activation. ^1^

**Fig. 1.**
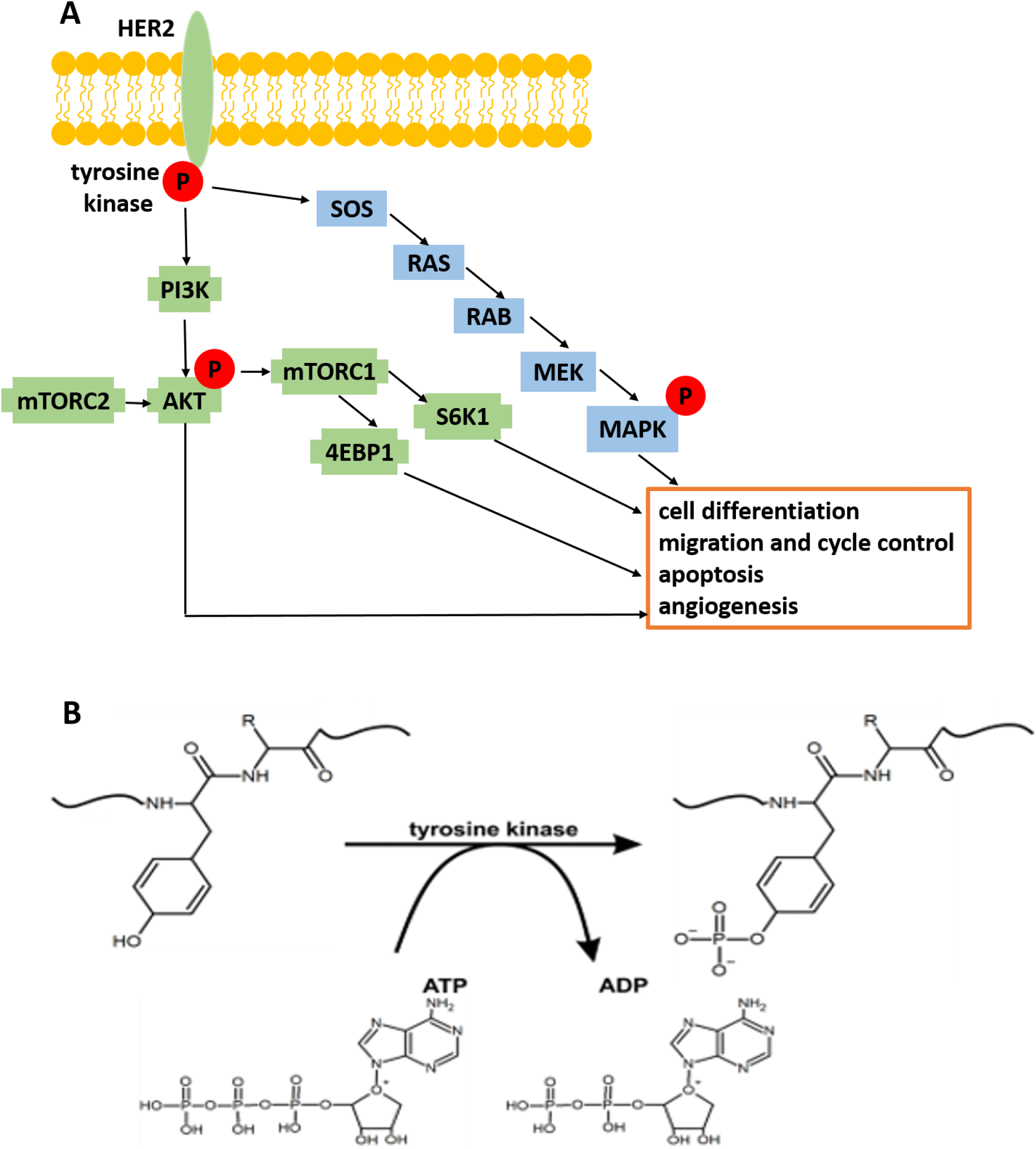
HER2 signaling pathways (panel A) the effect of phosphorylation catalyzed by tyrosine kinase (panel B).

Activated HER2 triggers phosphorylation of multiple tyrosine (Y) residues at its C-terminus (Fig.1B). Once activated RTKs play an important role in mediating cell-to-cell communication and controlling a wide range of complex biological functions, including cell growth, motility, differentiation, and metabolism by coupling to PI3K/Akt, Raf/MEK/MAPK, JAK/STAT, Src, PLCγ and other pathways. ^24^ (Fig.1A)

In normal cells, the activity of RTKs is governed by mechanisms keeping cell growth under control. Inappropriate activation of HER family receptors, overexpression of HER receptors signal and alterations of RTK pathways results in serious implications in cancer development leading to uncontrolled growth and spread of tumor cells. ^25^

Tyrosine phosphorylation is one of the leading areas of of biomedical research because of its relation to many human diseases including cancer via the dysregulation of receptor tyrosine kinases.

Protein phosphorylation belongs to the most important covalent post-translational modifications in cell signaling pathways catalysed by protein kinases, which represents the reversible attachment of a phosphate group to a protein.

Raman spectroscopy and imaging are unique tools that can monitor biochemical profile, localization and biocomposition alterations in specific organelles of cells as cancer develops. *In vitro* human cell line models have been widely used to predict clinical response to mechanisms associated with cancer development. In the current study, five breast cell lines: MCF-10A, MCF-7, MDA-MB-231, HTB-30 (SK-BR-3) and AU-565 were analysed.

MCF-10A human cells are used as a model for normal, non-tumorigenic epithelial human breast cells and represent normal level of HER2. The other studied cell lines represent HER2-positive or HER2-negative breast cancer models. When cells contain higher than normal levels of HER2 they are called HER2-positive. MCF-7 human cell line has an epithelial-like morphology and represents the human triple-positive breast carcinoma. Triple-positive breast cancer cells use HER2, estrogen receptors, progesterone receptors to grow. The HER2-positive cancers tend to grow and spread faster than HER2-negative breast cancers, but respond more effective to treatment with drugs that target the HER2 protein. MCF-7 is considered to be a poorly- aggressive and non-invasive cell line with low metastatic potential. In contrast, MDA-MB-231 is invasive, poorly differentiated and highly aggressive triple-negative breast cancer cell line without HER2 amplification as well as estrogen receptor (ER) and progesterone receptor (PR) expression.^26^ ^27^ Triple-negative breast cancers are usually more aggressive, more difficult to treat, and more likely to recur than cancers that are hormone receptor-positive or HER2- positive.

HTB-30 (SK-BR-3) line represents human adenocarcinoma that overexpresses the HER2 and is hormone-independent. ^28^ The AU-565 cell line overexpresses the HER2 as well as the HER- 3, HER-4 and p53 oncogenes.

Conventional assays used in clinical practice to identify HER2 status in breast cancer include immunohistochemistry (IHC) and in situ hybridization (ISH), both of which have limitations.^29^

In the current study, we studied HER2 expression in the human breast cancer cell lines as a model to explore the potential utility of a novel immunodetection technique based on Raman spectroscopy and Raman imaging combined with methods of artificial intelligence described in 30 31 32

## Results

Tyrosine phosphorylation is one of the most important post-translational modifications in cancer cells. It is very difficult to monitor these modifications at the genomic level because a large number of proteins (approximately 10,000) encoded by the human genome contain covalently bound phosphate. In proteomic approach , the situation is better because the typical protein kinase can attach phosphates to only 20 proteins ^33^. One of these proteins is cytochrome *c*. It has been proposed that reversible phosphorylation of cytochrome *c* mediated by cell signaling pathways is primary regulatory mechanism in living species that determines mitochondrial respiration, electron transport chain (ETC) flux, proton gradient ΔΨm, ATP production, and ROS generation. As it is well known these processes regulate efficiency of the oxidative phosphorylation and is directly related to many human diseases, including cancer, through a lack of energy, ROS production, cytochrome *c* release, and activation of apoptosis. ^34^ Because of the importance that phosphorylation has on biological processes in general, a huge emphasis has been focused on understanding the biological role of protein phosphorylation in human diseases. Current approaches of detection strategies include kinase activity assays, immunodetection, phosphoprotein or phosphopeptide enrichment. ^35^

We propose a novel method to monitor HER2 modifications at the molecular level via unique vibrational signatures of proteins.

The mechanism of tyrosine phosphorylation is presented in Figure 1B. The transfer of phosphate group to proteins is facilitated by enzymes called tyrosine kinases. The co-substrate for almost all protein kinases including tyrosine kinase is the donor of phosphate group called adenosine triphosphate (ATP). Tyrosine kinase facilitates the attack of a nucleophilic (–OH) group on tyrosine protein residue on the terminal phosphate group (γ-PO3^2-^) on ATP, resulting in the transfer of the phosphate group to tyrosine to form phosphotyrosine and ADP. Protein phosphorylation in a cell is a reversible, dynamic process that is mediated by kinases and phosphatases enzymes, which phosphorylate and dephosphorylate substrates, respectively.^33^ The structural formulae of tyrosine and phosphotyrosine are shown in Figure 2A.

**Fig. 2.**
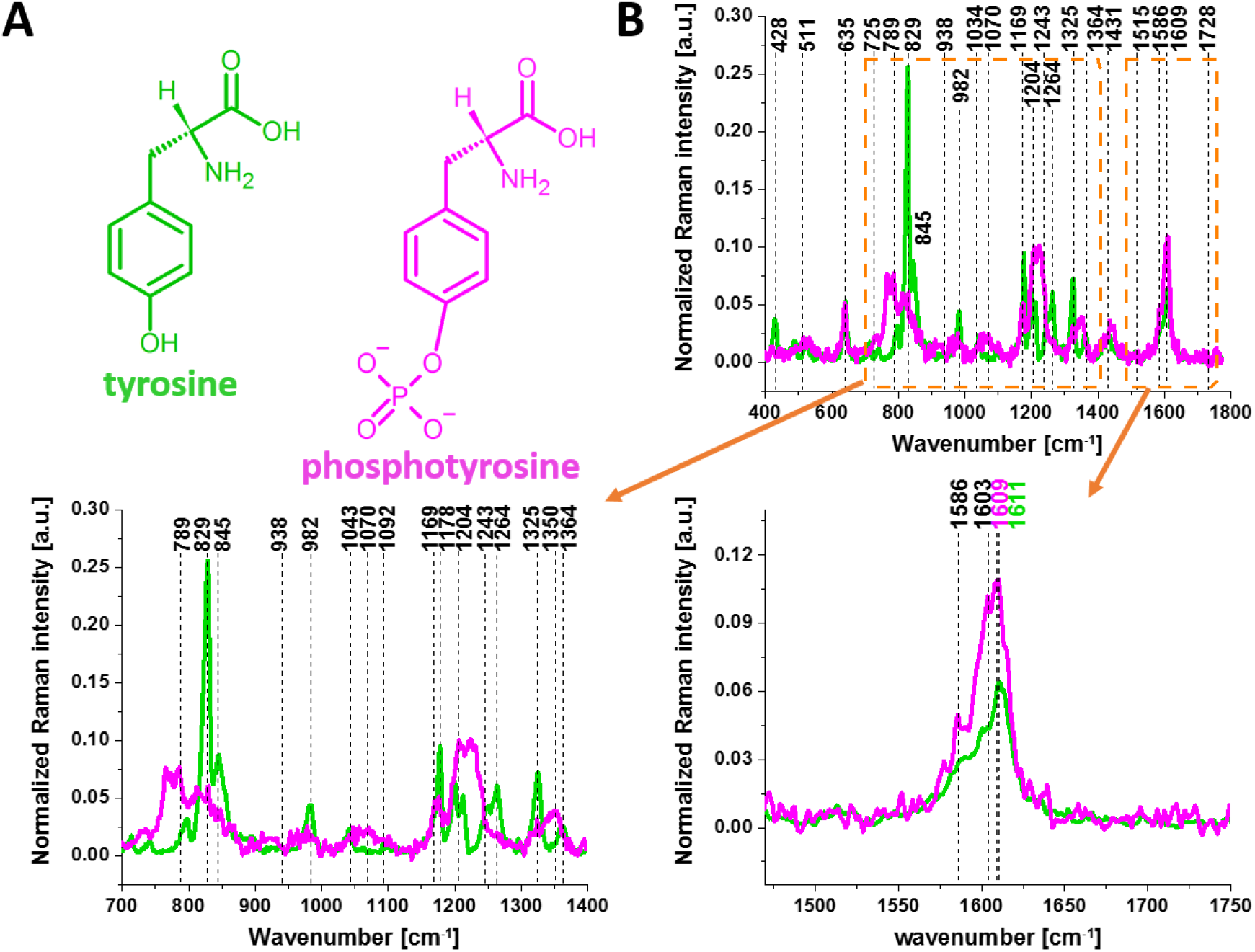
The structural formulae of tyrosine and phosphotyrosine (panel A) and Raman spectra of tyrosine (green line) and phosphotyrosine (magenta line) (panel B) 1

To study the activity of tyrosine kinase presented in Fig.1 by Raman spectroscopy and imaging let first concentrate on the Raman vibrational spectra of tyrosine and phosphorylated tyrosine (Fig.2B).

First, we concentrated on the changes in vibrational landscape that arise in tyrosine upon phosphorylation. The phosphorylation can be monitored by the spectral changes in proteins arising from either the phosphate stretching or amide vibrational modes.

The Raman spectra of tyrosine and phosphorylated tyrosine are shown in Fig. 2B. A detailed inspection into Figure 2B shows that tyrosine phosphorylation induces significant modifications in the Raman vibrational features: (1) an additional Raman peak at 1586 cm^−1^ close to the main peak at 1609 cm^−1^ that corresponds to the ring-O stretching mode ^36^ appears in phosphotyrosine; (2) the peak of phosphorylated tyrosine at 1609 cm^−1^ is shifted with respect to that of tyrosine observed at 1611 cm^−1^; (3) collapse into a single band upon tyrosine phosphorylation with a significant intensity decrease of the characteristic doublet (829 cm^−1^, 845 cm^−1^) of tyrosine corresponding to a Fermi resonance between the first overtone of the aromatic out-of-plane ring bend and the aromatic ring breathing fundamental ^37^; (4) the shift of the band at 1264 cm^−1^ corresponding to Amide III to 1243 cm^−1^ upon phosphorylation ^38^ ^39^.

The band positions of aromatic amino acids are sensitive to the microenvironment and may shift by up to 5 cm^−1^ in the Raman spectra of proteins ^40^. Vibrations of the PO4^−^ phosphate group of phosphorylated tyrosine are observed at 1070 cm^−1^ and corresponds to the O-P-O symmetric stretching mode ^37^ ^38^ ^39^ ^41^ ^42^ ^43^. The weak band at 1092 cm^−1^ band is due to the antisymmetric O-P-O stretching vibration ^39^ ^42^ ^43^. To summarize, most of the spectral shifts observed upon tyrosine phosphorylation are very similar to those observed in previously reported Raman studies. ^36^ ^40^ ^44^ ^45^ The band positions in the Raman spectra of the proteins compared to the reference tyrosine and phosphorylated tyrosine vary by up to only to few cm^−1^ due to their sensitivity to the microenvironment.

One of the important proteins sensitive to phosphorylation is cytochrome c. Figure 3 shows that the Raman spectra of cytochrome *c* in resonance with Q 0 -Q v electronic transitions are dominated by vibrational bands of the asymmetric A2g modes, i.e., 1585 cm ^−1^ (ν19), 1604 cm^−1^ (ν38 ) and from the B1g modes, i.e., 1638 cm ^−1^ (ν10 ), 1547 cm ^−1^ (ν11).^46, 47,48^. The band at 1585 cm^−1^ is primarily due to the methine bridge vibrations via Cα –Cm stretching and Cm –H bending modes respectively ^46^ ^49^ ^50^

**Fig. 3.**
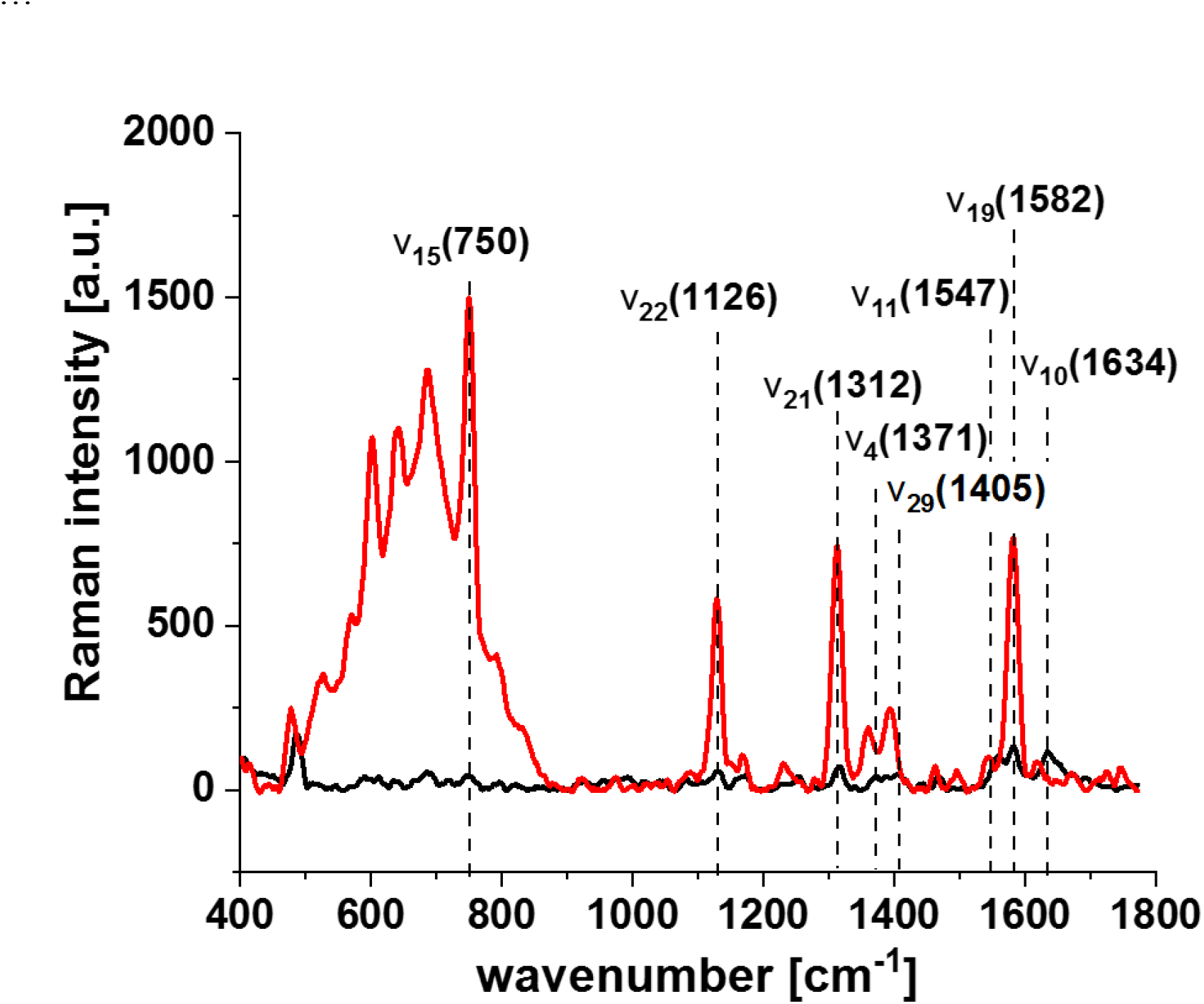
Raman spectra of isolated cytochrome c in solution (0.23 mM) dissolved in potassium phosphate buffer, pH 7.4, oxidized ferric Fe^3+^ (black line) and reduced ferrous Fe^2+^ (red line) cytochrome c. Ferrous cytochrome c was prepared by adding tenfold excess of reductor ascorbic acid.

The ν 15 vibration at 750 cm^-1^ is associated with the deformation vibrations of the 16 membered inner ring of heme group, ν4 vibration at 1371 cm^-1^ involves breathing- like motion of pyrrole ring , ν29 vibration at 1405 cm^-1^ represents antisymmetric stretching Ca-Cb, the ν19 is mainly due to methine bridge stretching Ca-Cm and Ca-Cb vibrations mixed with Cm-H bending mode with a perpendicular displacement of the Cm atom to the plane of the heme group ^49^…

Having obtained the reference Raman fingerprint of tyrosine phosphorylation and cytochrome *c*, we focused on the Raman vibrational modifications arising in the proteins due to tyrosine phosphorylation in human normal and cancerous breast *in vitro* cells.

Figure 4 shows the Raman images and Raman spectra for a typical cell of HTB-30 (SK-BR-3) line. These cells represent human adenocarcinoma and overexpresses the HER2/c-erb-2 gene product.

**Fig. 4.**
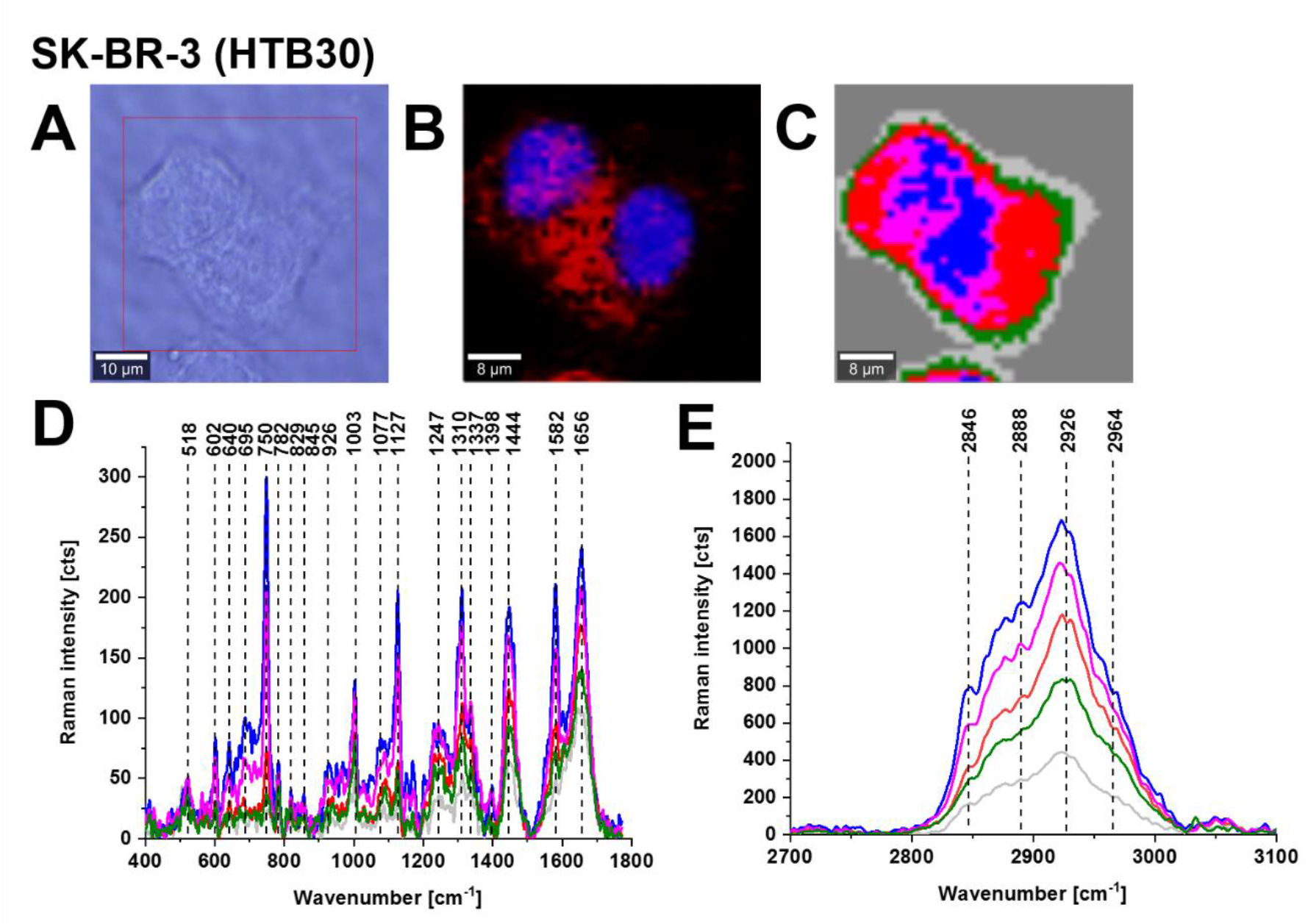
Raman imaging of a typical breast cancer cell (HTB-30). Microscopy images (**A**), fluorescence image of ER/lipid droplets—red (Oil Red O) (**B**), fluorescence image of nucleus-blue (Hoechst 33342 (**B**), Raman image (**C**) obtained from the Cluster Analysis (nucleus (red), endoplasmic reticulum (blue), lipid droplets (orange) cytoplasm (green), mitochondria (magenta), membrane (light grey), area outside the cell (dark grey), Raman spectra of the respective organelles (the same colour as in Raman image) in the fingerprint region **(D**) and high-frequency region (**E),** Resolution of Raman images 1 μm, integration time 0.3 s, 10 mW at 532 nm excitation and resolution of 1 μm in fluorescence images, integration time 0.01 s The colors of spectra correspond to the colors of cluster classes in the Raman maps.

Fig. 4 shows that the Raman spectrum of cancer breast cell of HTB-30 (SK-BR-3) line is dominated by cytochrome *c*. One can see that the vibrations of the heme group in cytochrome *c*: ν15 (750 cm^-1^), ν22 (1127 cm^-1^), ν21 (1310 cm^-1^) and ν19 (1582 cm^-1^) are the strongest bands in the of cancer breast cell of HTB-30 (SK-BR-3).

Detailed inspection into Fig. 5 shows (A) the characteristic doublet (835 cm^−1^, 846 cm^−1^) of tyrosine corresponding to a Fermi resonance between the first overtone of the aromatic out-of-plane ring bend and the aromatic ring breathing fundamental ^49^. The bands do not collapse into a single band as it happens upon tyrosine phosphorylation with a significant intensity decrease as can be seen in Fig. 1; (B) the band at 1270 cm^−1^ corresponding to Amide III is partially shifted to 1241 cm^−1^ upon phosphorylation ^38^ ^39^. The vibrational features of tyrosine in the cancerous HTB-30 (SK-BR-3) cells illustrate dynamic nature of phosphorylated proteins in a cell ^33^ and suggest that the equilibrium between tyrosine and phosphorylated tyrosine is shifted towards non-phosphorylated tyrosine in breast cancer cells.

**Fig. 5.**
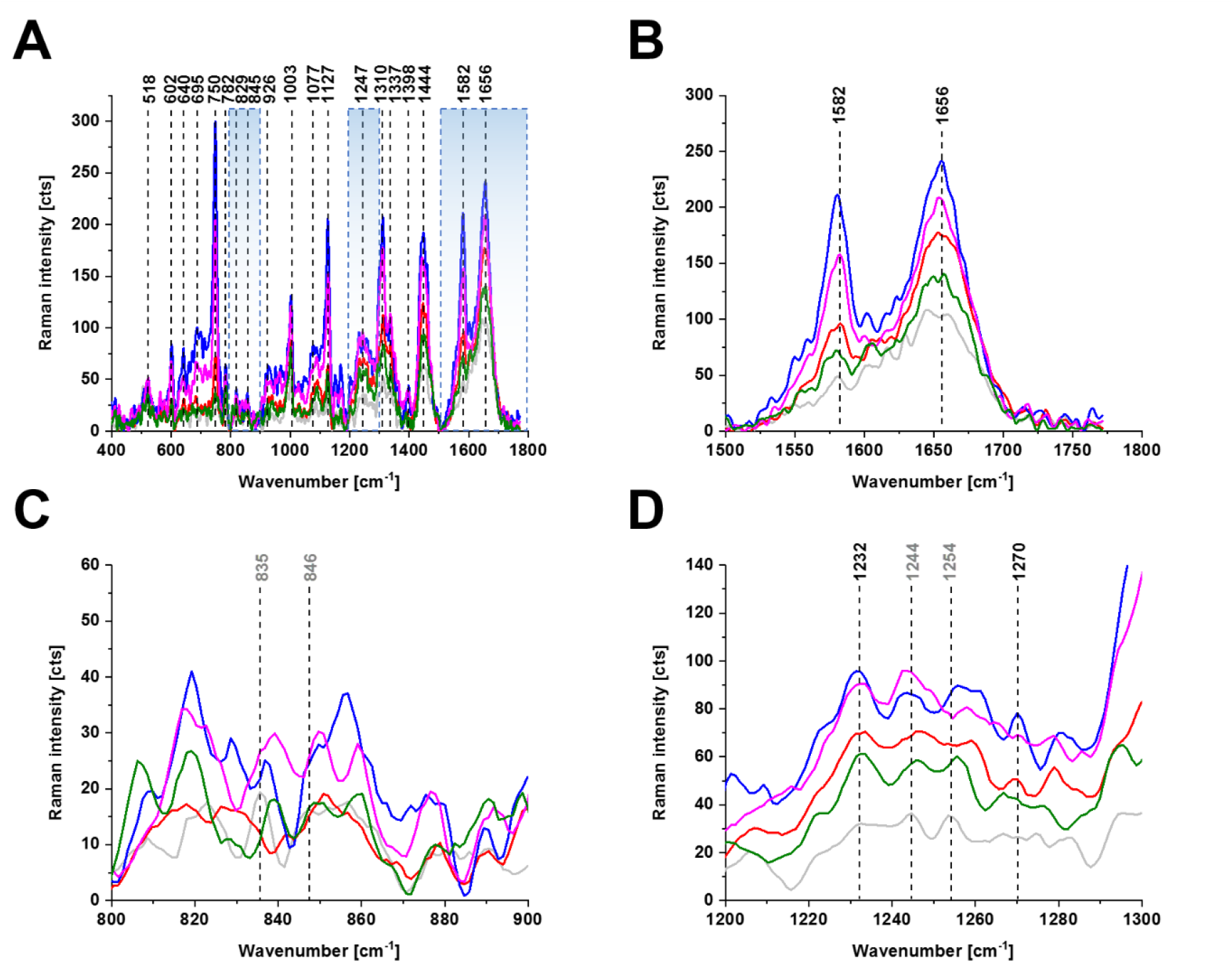
Raman spectra of a typical breast cancer cell (HTB-30) for the cell organelles (nucleus (red), endoplasmic reticulum (blue), lipid droplets (orange), cytoplasm (green), mitochondria (magenta), membrane (light grey).

Fig. 5C shows the Raman bands of cytochrome *c* at 1582 cm^-1^ and amide I at 1656 cm^-1^ of a typical breast cancer cell (HTB-30). The equilibrium between the ferric Fe^3+^ and ferrous Fe^2+^ forms is evidently shifted towards ferrous cytochrome *c* in breast cancer cells that overexpress the HER2 proteins.

In the current study, six breast cancer cell lines: MCF-10A, MCF-7, MDA-MB-231, HTB-30 (SK-BR-3) and AU-565 were studied and the vibrational assignments of the Raman bands at 532 nm laser excitation are shown in Table 1.

**Table 1.** The vibrational assignments of the Raman bands observed in MCF-10A, MCF-7, MDA-MB-231, HTB-30 (SK-BR-3) and AU-565 at 532 nm laser excitation.

| HTB-30<br>HER-2/neu<br>oncogene<br>adenocarcin<br>oma | MCF-10A<br>Control<br>normal | MCF-7<br>Triple -<br>positive<br>poorly-<br>aggressive | MDA-MB-231<br>Triple-negative<br>Aggressive<br>cancer | AU-565<br>HER-2/neu<br>oncogene<br>adenocarcino<br>ma | Vibrational<br>assignments<sup>47 51</sup> |
| --- | --- | --- | --- | --- | --- |
| 483 | 483 | 483 | 485 | 479 | glycogen |
| 518 | 518 | 518 | 520 | 521 | phosphatidylinositol |
| 602 | 602 | 602 | 602 | 601 | phosphatidylinositol |
| 640 | 640 | 641 | 638 | 641 | C-S stretching and<br>C-C twisting of<br>proteins-<br>tyrosine |
|  |  |  |  |  | C-C twisting mode of tyrosine |
| 689 | 689 | 689 | 687 | - | cytochrome *c* |
| 726 | 726 | 726 | 726 | - | C-S (protein), CH<sub>2</sub> rocking, adenine |
| 748 | 749 | 746 | 746 | 748 | *c*, *cI* and *b*-types of cytochrome<br>ν<sub>15</sub> deformation of inner ring of heme group |
| 782 | 785-787 | 783 | 785 | 785 | DNA: O-P-O, cytosine, uracil, thymine<br>Phosphatidylserine |
| 829 | 829 | 826 | 826 | 823 | tyrosine (Fermi resonance of ring fundamental and overtone) |
| 853 | 845 | 848 | 849 | 852 | tyrosine (Fermi resonance of ring fundamental and overtone) |
| 877 | 877 | 877 | 877 | - | C-C-N<sup>+</sup> symmetric stretching (lipids choline group) |
| 932 | 932 | 936 | 935 | 932 | skeletal C-C, α-helix |
| 1002 | 1002 | 1001 | 1001 | 1003 | phenylalanine |
| 1085 | 1080-1085 | 1084 | 1085 | 1080<br>1092 | C-C (lipid)<br>Symmetric phosphate stretching vibration of ν<sub>3</sub>PO<sub>4</sub> |
| 1126 | 1124 | 1126 | 1125 | 1126 | *c*, *cI* and *b*-types of cytochrome |
| 1169 | 1169 | 1169 | 1168 | 1171 | tyrosine<br>δ(C-H), tyrosine (protein assignment) |
| 1248 | 1241 | 1247 | 1248 | 1246 | phosphorylated protein Amide III |
| 1264 | 1254 | 1263 | 1258/1265 | 1258 | amide III |
| 1311 | 1306 | 1306 | 1302 | 1308 | C -type cytochromes, CH<sub>3</sub>CH<sub>2</sub> twisting mode of collagen/lipid |
| 1336 | 1336 | 1335 | 1335 | 1337 | cytochrome *b* |
| 1444 | 1444 | 1440 | 1436 | 1436 | Cholesterol, CH<sub>2</sub> deformation |
| 1446 | 1446 | 1446 | 1445 | 1447 | CH<sub>2</sub> bending mode of proteins and lipids<br>CH<sub>2</sub> deformation |
| 1456 | 1456 | 1456 | 1459 | 1460 | CH<sub>2</sub> stretching/CH<sub>3</sub> asymmetric deformation overlapping asymmetric CH<sub>3</sub> bending and CH<sub>2</sub> scissoring (is associated with collagen, & phospholipids) |
| 1578 | 1576 | 1578 | 1578 | 1576 | pyrimidine ring (nucleic acids) and heme protein |
| 1582 | 1582 | 1582 | 1582 | 1582 | cytochrome c v<sub>19</sub> methine bridge vibrations via C<sub>α</sub>–C<sub>m</sub> stretching and C<sub>m</sub>–H bending modes |
| 1602 | 1600 | 1602 | 1602 | 1602 | C=C in-plane bending mode of phenylalanine and tyrosine |
| 1617 | 1618 | 1617 | 1614 | 1618 | C=C phenylalanine, tyrosine |
| 1634 | 1631 | 1633 | 1632 | 1632 | amide I |
| 1647 | - | 1647 | 1648 | 1647 | random coils |
| 1656 | 1655 | 1655 | 1653 | 1654 | amide I |
| 2846 | 2846 | 2846 | 2845 | 2846 | CH<sub>2</sub> symmetric stretch of lipids |
| 2888 | 2884 | 2884 | 2890 | 2878 | CH<sub>2</sub> asymmetric stretch of lipids and proteins |
| 2926 | 2929 | 2929 | 2926 | 2926 | symmetric CH<sub>3</sub> stretch due primarily to protein |
| 2964 | 2964 | 2964 | 2952 | 2964 | v<sub>as</sub> CH<sub>3</sub>, lipids, fatty cholesterol and cholesterol ester |
| 3012 | 3012 | 3012 | 3012 | 3012 | unsaturated =CH stretch |
| 3055 | 3055 | - | - | 3054 | aromatic ring =CH, tyrosine |

In Fig. 7 we showed average normalized Raman spectra of breast cancer cells (HTB-30) (A), AU-565 (B) and MCF-7 (C) for the cell organelles.

**Fig. 6.**
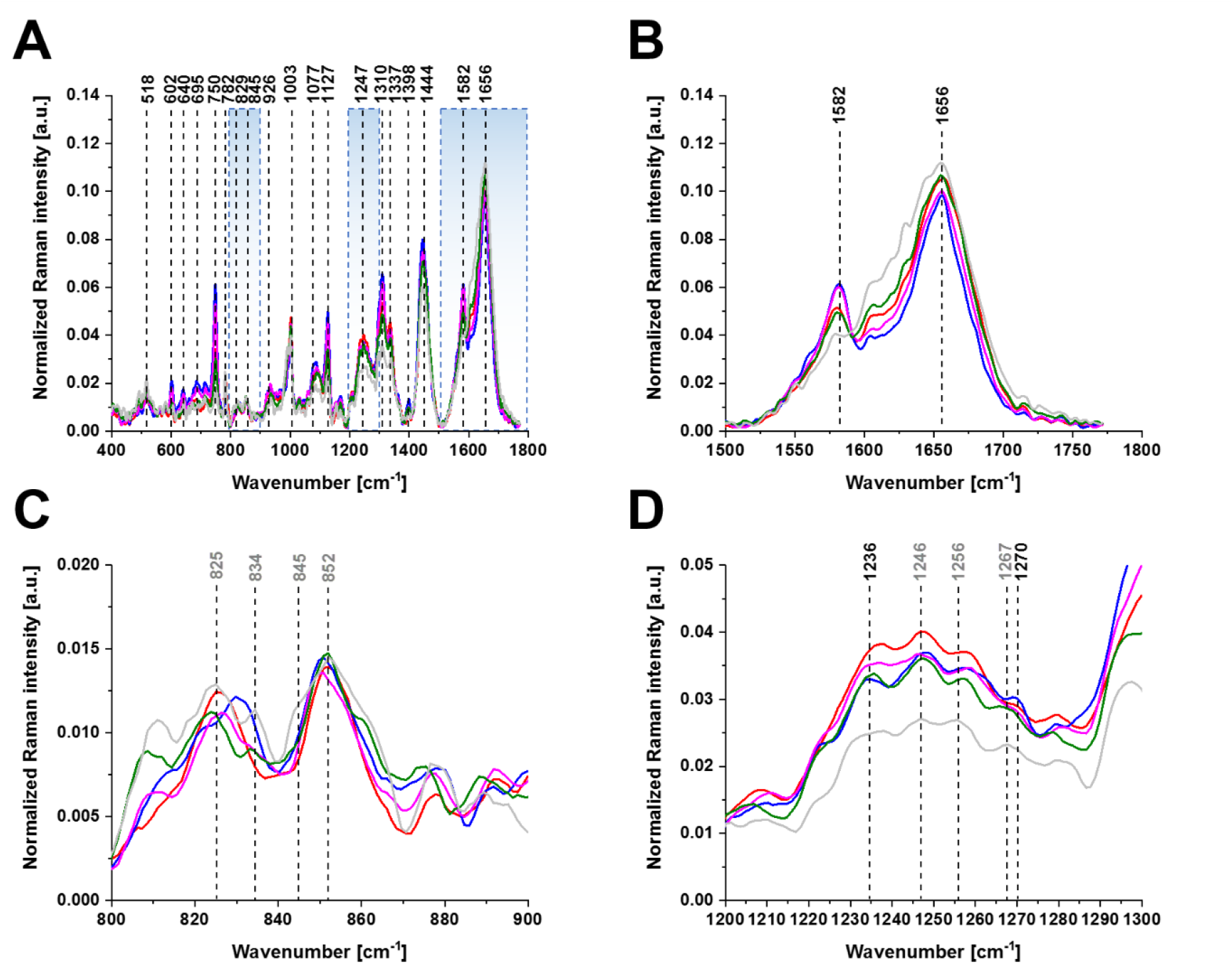
Average normalized Raman spectra of breast cancer cells (HTB-30) for the cell organelles (nucleus (red), endoplasmic reticulum (blue), lipid droplets (orange) cytoplasm (green), mitochondria (magenta), membrane (light grey), average Raman spectra were obtained for the number of cells, n=9.

**Fig. 7.**
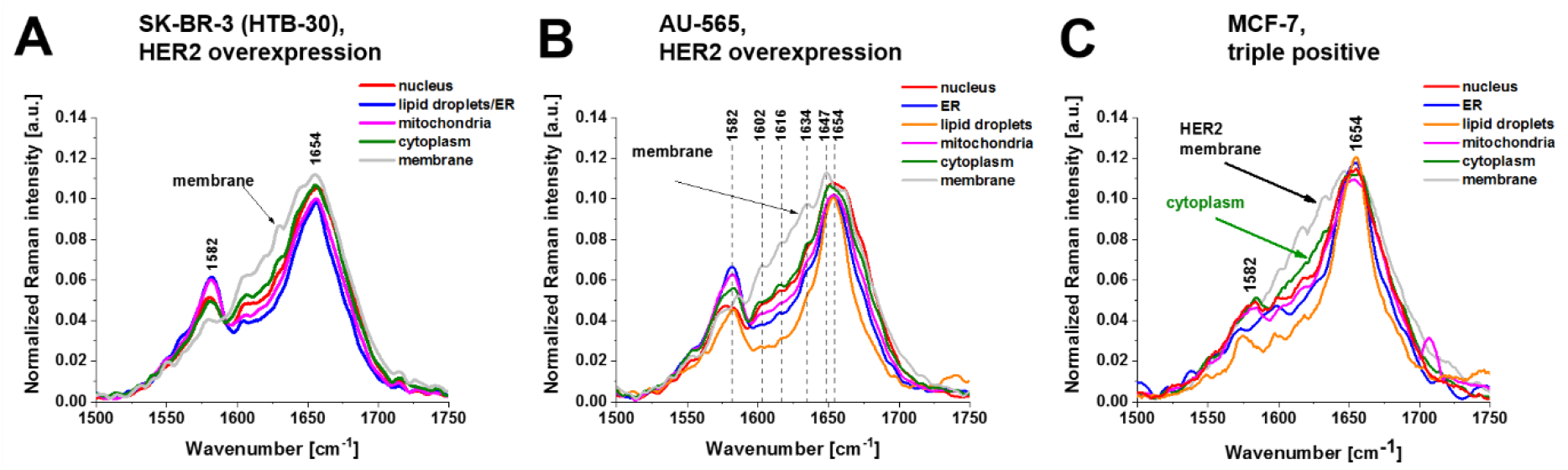
Average normalized Raman spectra of breast cancer cells (HTB-30) (A), AU-565 (B) and MCF- 7 (C) for the cell organelles (nucleus (red), endoplasmic reticulum (blue), lipid droplets (orange) cytoplasm (green), mitochondria (magenta), membrane (light grey).

Fig. 7 shows average normalized Raman spectra of breast cancer cells (HTB-30) (A), AU-565 (B) and MCF-7 (C) for the cell organelles. These three cancer cell lines overexpress HER2 on the surface of cells. Detail inspection into Fig. 7 shows that the vibrational profiles of cytochrome *c* at 1582 cm^-1^ and protein Amide I profile at 1654 cm^-1^ in the cell membranes differ spectacularly from the other organelles. Additional bands at 1602 cm^-1^, 1618 cm^-1^, 1634 cm^-1^, 1647 cm^-1^ appear (Table 1) and most of them correspond to tyrosine vibrations. The presented in Fig. 7 cells have HER2 protein expression on the surface of the cell that belongs to a family of receptor tyrosine kinases (RTKs) and consists of an extracellular domain that includes four subdomains (I-IV) [16], a single helix transmembrane lipophilic segment and an intracellular region that contains a tyrosine kinase domain (TKD).

Our results suggests that we found a novel detection methodology for HER2 protein quantitation in breast cancer cells using Raman spectroscopy and Raman imaging.

The conventional methods of immunohistochemistry give a score of 0 to 3+ that measures the amount of HER2 proteins on the surface of cells in a breast cancer tissue sample. If the score is 0 to 1+, it’s considered HER2-negative. If the score is 2+, it’s considered borderline. A score of 3+ is considered HER2-positive.

To confirm that the HER2 status can be determined by Raman spectroscopy we extended the list of studied cancer breast cell lines to the normal cells (MCF-10A) and triple-negative aggressive breast cancer (MDA-MB-231).

Fig. 8 A shows average normalized Raman spectra of membranes in breast cancer cells: triple-positive MCF-7, HTB-30 and AU-565 overexpressing HER2, the normal cells (MCF-10A) (HER2 at the normal level) and triple-negative aggressive breast cancer (MDA-MB-231).

**Fig. 8.**
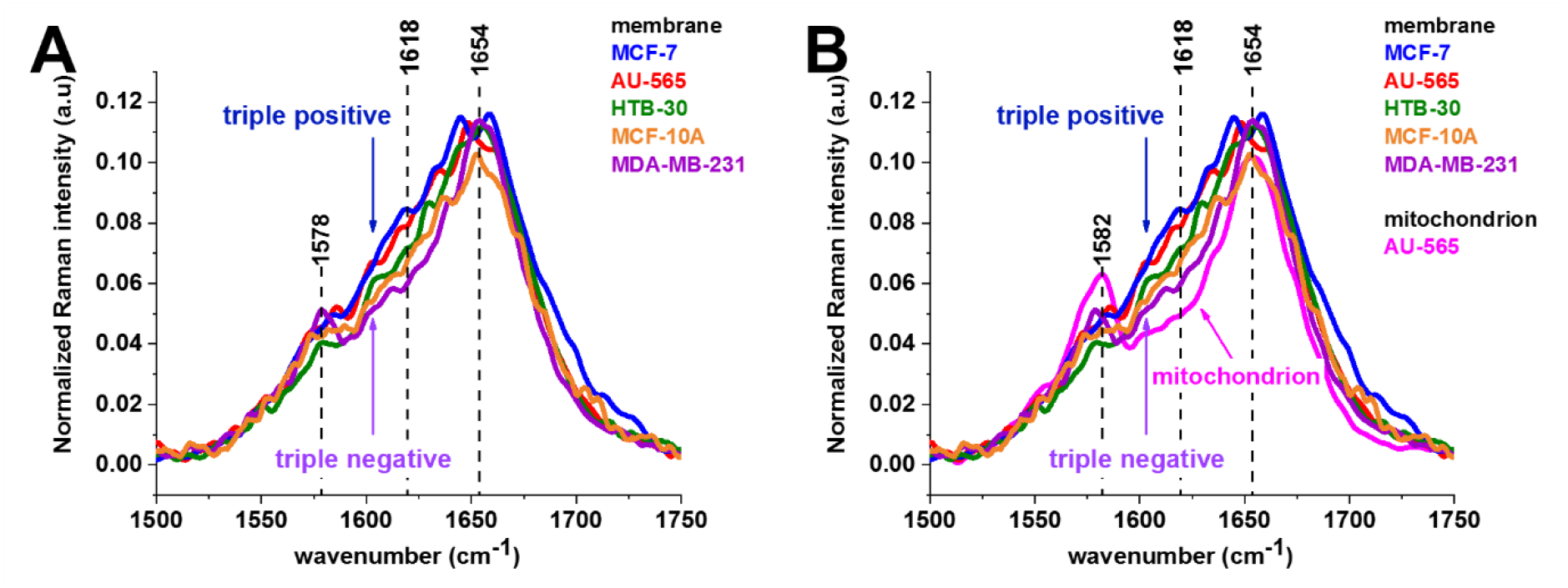
Average normalized Raman spectra of membranes (A, B) and mitochondria (B) in breast cancer cells, triple-positive MCF-7, HTB-30 and AU-565 overexpressing HER2, the normal cells MCF-10A (HER2 at the normal level) and triple-negative aggressive breast cancer MDA-MB-231.

Fig. 8 B shows average normalized Raman spectra of membranes and mitochondria in breast cancer cells (HTB-30) (A), triple-positive HER2 MCF-7 (B), HTB-30 and AU-565 (C) overexpressing HER2, the normal cells (MCF-10A) (HER2 at the normal level) and triple - negative aggressive breast cancer (MDA-MB-231).

Fig. 8 B shows average normalized Raman spectra of membranes and mitochondria in breast cancer cells, triple-positive HER2 MCF-7, (HTB-30) and AU-565 (C) overexpressing HER2, the normal cells (MCF-10A) (HER2 at the normal level) and triple -negative aggressive breast cancer (MDA-MB-231). One can see that that the highest concentration of cytochrome *c* represented by the vibration at 1584 cm^-1^ is located in mitochondria and support the results reported recently^1^

Detailed inspection into Fig. 8B shows that the Raman intensity between the cytochrome *c* band at 1582 cm^-1^ and the protein amide I at 1654 cm^-1^ is spectacularly different for mitochondria and membranes of the studied cells. The bands in this region correspond to tyrosine kinase vibrations (Table 1). We have chosen for analysis the Raman intensity at 1618 cm^-1^ representing tyrosine kinase of HER2. The results for the concentration of tyrosine kinase illustrated by the Raman intensity at 1618 cm^-1^ are presented in Fig. 9. One can see that the Raman intensity proportional to concentration of tyrosine kinase receptor at the intracellular side of the membrane for triple-positive MCF-7, HTB-30 and AU-565 overexpressing HER2 is higher than that for the normal cells (MCF-10A) (HER2 at the normal level). In contrast the triple-negative aggressive breast cancer (MDA-MB-231) exhibits lower Raman signal than the cells (MCF- 10A) with HER2 at the normal level.

**Fig. 9.**
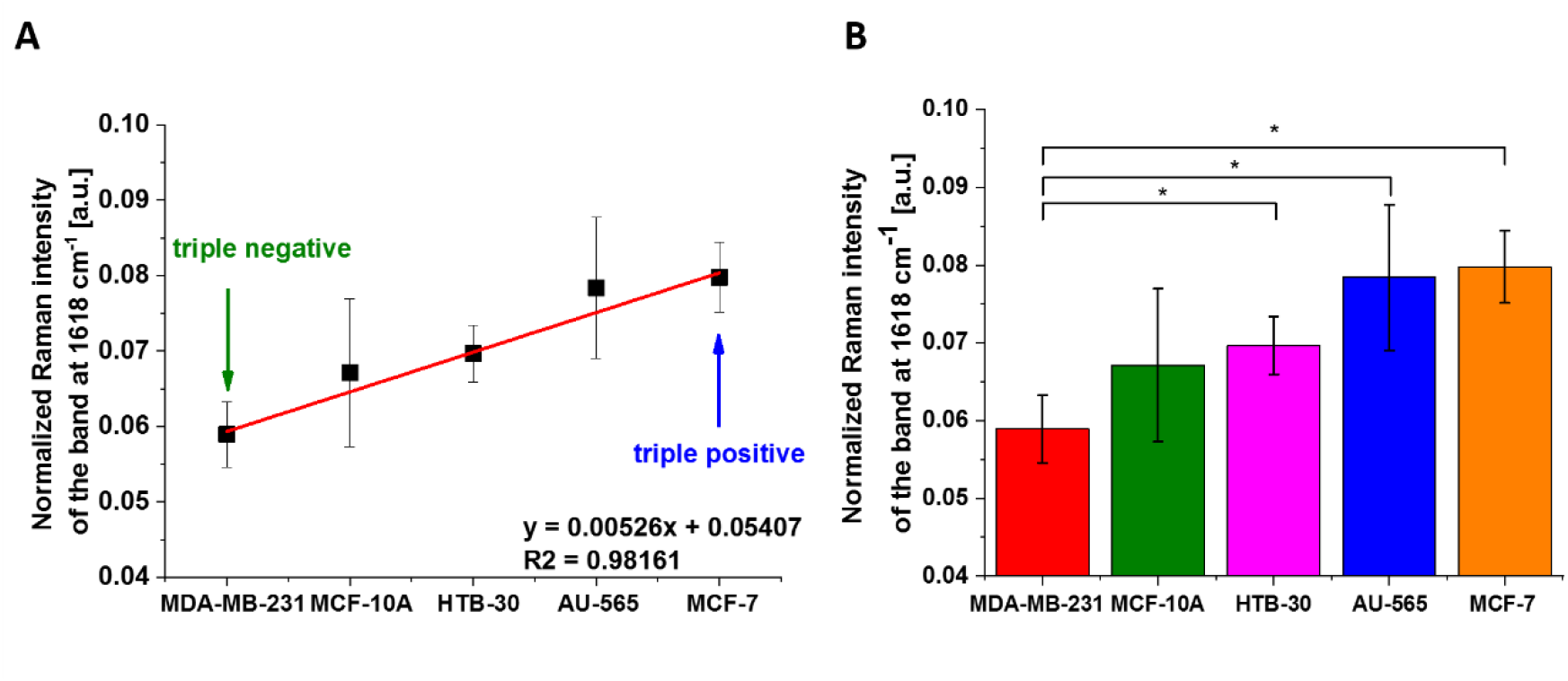
Normalized Raman intensity of the band at 1618 cm^-1^ of breast cancer cells membranes for triple-positive (MCF-7), overexpressing HER2 (HTB-30 and AU-565), the normal cells (MCF-10A) (HER2 at the normal level) and triple - negative aggressive breast cancer (MDA-MB-231), (A) linear fitting R^2^=0.98161. (B) normalized Raman band intensity (1618 cm^-1^) as a function of breast cancer (mean ± SD); the statistically significant results have been marked with an asterix (p < 0.05).

Fig. 9 shows average normalized Raman intensity at 1618 cm^-1^ at membranes (A,B) and mitochondria (B) in breast cancer cells: triple-positive MCF-7 (B), (HTB-30) and AU-565 (C) overexpressing HER2, the normal cells (MCF-10A) (HER2 at the normal level) and triple - negative aggressive breast cancer (MDA-MB-231). Fig. 9 shows the linear dependence of the normalized Raman intensity at 1618 cm^-1^ of tyrosine vibration status on the surface of the normal and cancer cells vs. on the HER2 and EGFR status. The HER2 expression in these cells correlates in an excellent way with the conventional biological methods of HER2 determination. ^28^

One can see that Raman spectroscopy shows strong correlations with the results of conventional HER2 testing methodologies by IHC analysis presented in Table 2 for all studied breast cancer cell lines. In contrast, there is no correlation with EGFR receptors. Our results for HER2 receptor kinases in breast cancer are supported by other reports in the literature that the dimerization causes the activation *of* downstream signaling pathways by phosphorylating the intracellular tyrosine kinase domain. ^52^

**Table 2.**
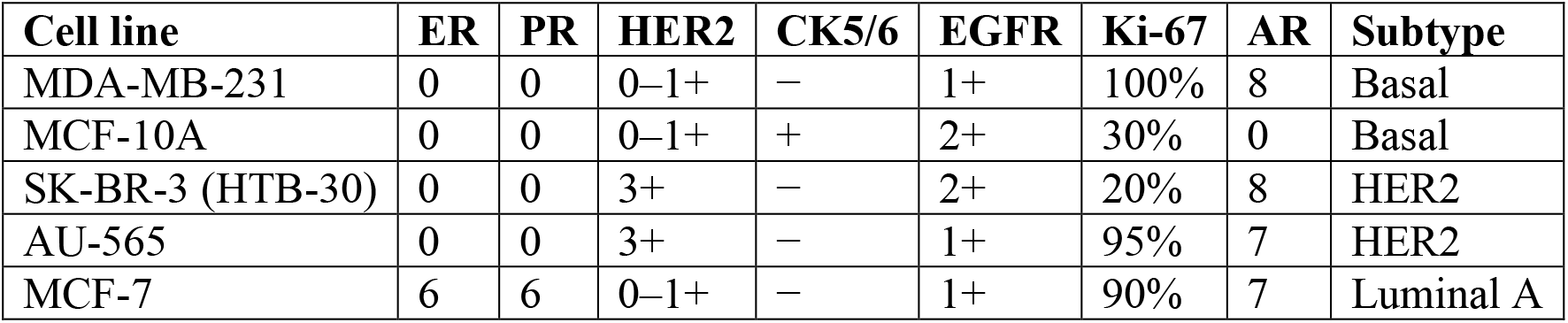
The results of IHC analysis for breast cancer cell lines.^57^

No ligands for HER2 have yet been identified yet. ^21^ ^22^ and dimerization with any of the other three subdomains is considered to activate HER2. ^23^ The dimerization in the extracellular region of HER2 induces intracellular conformational changes that trigger tyrosine kinase activation. ^1^

The intriguing exception is MCF-7 cancer which does not correlate with HER2 (0-1+) but ER and PRare very high, being 6, 6 respectively. It was reported that both estrogen and progesterone can interact with membrane receptors. ^53^

Breast cancers are categorized into five different subtypes, luminal A/B, HER2-positive (HER^+^), basal-like, claudin-low, and normal breast-like, based on the expression levels of estrogen and progesterone receptors (ER and PR), HER2, cytokeratins 5/6, and claudins 3/4/7. 54 55 56

Fig. 10 shows the average normalized Raman intensity of cytochrome *c* at 1582 cm^-1^ in mitochondria of breast cancer cells: triple-positive HER2 MCF-7 (B), HTB-30 and AU-565 (C) overexpressing HER2, the normal cells MCF-10A (HER2 at the normal level) and triple-negative aggressive breast cancer MDA-MB-231.

**Fig. 10.**
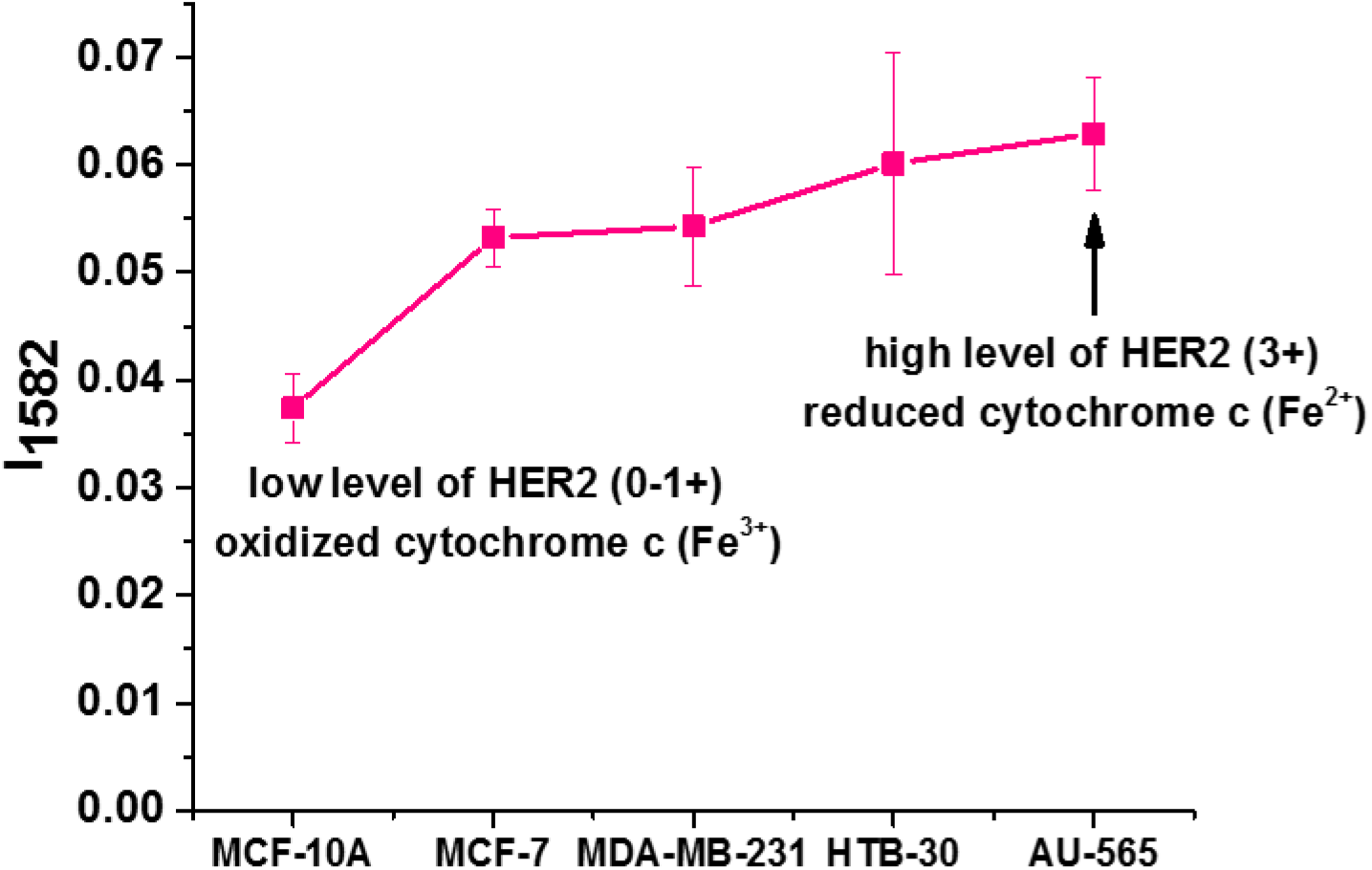
Average normalized Raman intensity of cytochrome c at 1582 cm^-1^ in mitochondria of breast cancer cells: triple-positive HER2 MCF-7, HTB-30 and AU-565 overexpressing HER2, the normal cells MCF-10A (HER2 at the normal level) and triple-negative aggressive breast cancer MDA-MB-231.

One can see that the Raman intensity of cytochrome *c* shows a strong and statistically significant correlation with HER2. The increase in the Raman signal results from a change in the redox status from the oxidized Fe^3+^ to the reduced form Fe^2+^. ^4^ ^58^ For cells expressing the high level of HER2 cytochrome *c* in mitochondria exhibits significant shift towards reduced form Fe^2+^.

## Conclusions

*In vitro* human cell lines are one of the most commonly used preclinical models to the biology of cancer to examine the factors influencing the response of tumors to therapy. This paper demonstrates that Raman spectroscopy and Raman imaging have great potential for the quantitative identification of proteins in routine clinical tests. We have shown that Raman approach demonstrate sensitive and rapid quantitation of HER2 protein in five breast cell lines: MCF-10A, MCF-7, MDA-MB-231, HTB-30 (SK-BR-3), AU-565. The results obtained from this methodology are highly correlated with routine clinical HER2 testing in *in vitro* human cell line breast cancer models and are very promising for predicting pathologic response to drug therapy. Raman spectroscopy shows strong correlations with the results of routine HER2 testing methodologies by IHC analysis. The Raman intensity of cytochrome *c* at 1582 cm^-1^ shows a strong and statistically significant correlation with HER2. The increase in the Raman signal results from shift in the redox status from the oxidized Fe^3+^ to the reduced form Fe^2+^. We showed that for cells expressing the high level of HER2 at the membrane cytochrome *c* in mitochondria exhibits a significant shift towards reduced form Fe^2+^. The intriguing exception is MCF-7 cancer for which Raman measurements do not correlate with HER2 (0-1+) but correlate with ER and PR receptors expression Our findings should be considered as preliminary until they will be confirmed by larger than five, well-controlled *in vivo* cell lines and *ex vivo* tissues of patients with HER2-positive and negative breast cancer. Our data suggest that the quantitative Raman measurement of HER2 protein levels may offer an advantage over current clinical testing and will help in proper selection of patients for HER2-targeted therapy. Further studies in *ex vivo* tissues for large patient cohort are warranted.

## Materials and Methods

### Reference Chemicals

Cytochrome *c* (no. C2506), tyrosine (no. T3754), phosphotyrosine (no. P9405) were purchased from Merck Life Science.

### Cell Cultures Preparation for Raman measurements

Cell lines MCF10A (no. CRL-10317, ATCC), MCF7 (no. HTB-22, ATCC), MDA-MB-231 (no. CRM-HTB-26, ATCC), SK-BR-3 (no. HTB30, ATCC) and AU565 (no. CRL-2351, ATCC) were persuaded from ATCC and culture according to ATCC protocols. For Raman measurements cells were seeded onto calcium fluoride windows (CaF2, 25 × 2 mm) at a low density of 5 × 10^4^. After 24 h, the CaF2 slides with cells were rinsed with phosphate-buffered saline (PBS, SIGMA P-5368, pH 7.4 at 25 °C, c = 0.01 M) to remove any residual medium, then cells were fixed with 4% formalin solution (neutrally buffered) and kept in PBS (no. 10010023, Gibco) during the experiment. After performing Raman imaging measurements, the cells were treated by 15 minutes with Hoechst 33342 (25 μL at 1 μg/mL per mL of PBS) and Oil Red O (10 μL of 0.5 mM Oil Red dissolved in 60% isopropanol/dH2O per each mL of PBS) Following a PBS wash, the cells were fluorescence imaged using an Alpha 300RSA WITec microscope, with the addition of fresh PBS.

### Raman Spectroscopy and Imaging

Raman measurements of the human breast cell lines were performed with a WITec confocal alpha 300 Raman microscope and a diode laser coupled to the microscope via an optical fiber of 50 μm diameter core. The 40× objective (NIKON CFI Plan Fluor C ELWD (Extra-Long Working Distance) 40×: N.A. 0.60, W.D. 3.6–2.8 mm; DIC-M, C.C.0-2) was used. Before collection of Raman spectra a standard single-point calibration procedure was performed with the reference of Raman band produced by a silicon plate at 520.7 cm^−1^. The Raman spectra were induced by a 532 nm excitation wavelength laser with the power of 10 mW in the focus spot and an integration time of 0.3 s by Andor Newton DU970-UVB-353 CCD camera in enhanced mode (EMCCD). The Raman data analysis was performed using WITec (WITec Project Plus 4) and OriginPro 2023 software. To construct the Raman maps presented in the paper we used Cluster Analysis procedure described in detail in ^30^ ^31^ ^32^. The number of clusters used for analysis was 6 (the minimum number of clusters characterized by different average Raman spectra, which describe the organelles in the cell: nucleus, lipid droplets, endoplasmic reticulum (ER), cytoplasm, mitochondria, cell border). The colors of the cluster images correspond to the colors of the Raman spectra of nucleus (red), lipid droplets (orange), endoplasmic reticulum (blue), cytoplasm (green), mitochondria (magenta), and cell border (light grey). Number of analyzed cells n(MCF-7)=4, n(HTB-30)=11, n(AU-565)=20, n(MDA-MB-231)=3; n(MCF-10A)=3 number of Raman spectra of MCF-7, HTB-30, AU-565, MDA-MB-231, MCF- 10A used for averaging 7424, 18125, 8639, 10920, 5200, respectively.

## ANOVA

The one-way ANOVA test implemented in OriginPro 2016 software was applied for the statistical analysis of the spectroscopic data. To calculate the value of statistical significance the Tukey test was used; asterisk * denotes that the differences are statistically significant, p-value ≤ 0.05.

## Author Contributions

Conceptualization: H.A.; Funding acquisition: H.A.; Investigation: J.M.S., M.K.; Conducted a series of laboratory experiments and analyzed data: J.M.S, M.K., H.A.; Methodology: J.M.S., M.K. and H.A.; Manuscript writing and editing: H.A., M.K., J.M.S. All authors have read and agreed to the published version of the manuscript.

## Data Availability Statement

The raw data underlying the results presented in the study are available from Lodz University of Technology Institutional Data Access. Request for access to those data should be addressed to the corresponding author.

## References

1. Abramczyk H, Imiela A, Brożek-Płuska B, Kopeć M, Surmacki J, Śliwińska A. Aberrant protein phosphorylation in cancer by using raman biomarkers. Cancers (Basel). 2019;11(12). doi:10.3390/cancers11122017

2. Kopec M, Imiela A, Abramczyk H. Monitoring glycosylation metabolism in brain and breast cancer by Raman imaging. Sci Rep. 2019;9(1):166. doi:10.1038/s41598-018-36622-7

3. Kopec M, Błaszczyk M, Radek M, Abramczyk H. Raman imaging and statistical methods for analysis various type of human brain tumors and breast cancers. Spectrochim Acta Part A Mol Biomol Spectrosc. 2021;262:120091. 10.1016/j.saa.2021.120091

4. Surmacki JM, Abramczyk H. Confocal Raman imaging reveals the impact of retinoids on human breast cancer via monitoring the redox status of cytochrome c. Sci Rep. 2023;13(1):1–11. doi:10.1038/s41598-023-42301-z

5. Brozek-Pluska B, Musial J, Kordek R, Abramczyk H. Analysis of Human Colon by Raman Spectroscopy and Imaging-Elucidation of Biochemical Changes in Carcinogenesis. Int J Mol Sci. 2019;20(14). doi:10.3390/ijms20143398

6. Slamon DJ, Godolphin W, Jones LA, et al. HER-2/neu Proto-oncogene in Human Bresat and Ovarian Cancer. Science (80- ). 1989; 244:707-712.

7. Slamon DJ, Clark GM, Wong SG, Levin WJ, Ullrich A, Mcguire WL. Human Breast Cancer: Correlation of Relapse and Survival with Amplification of.

8. Gutierrez C, Schiff R. HER2: biology, detection, and clinical implications. Arch Pathol Lab Med. 2011;135(1):55–62. doi:10.5858/2010-0454-RAR.1

9. Hynes NE, Stern DF. The biology of erbB-2/nue/HER-2 and its role in cancer. Biochim Biophys Acta - Rev Cancer. 1994;1198(2):165–184. 10.1016/0304-419X(94)90012-4

10. Klapper LN, Kirschbaum MH, Seta M, Yarden Y. Biochemical and Clinical Implications of the ErbB/HER Signaling Network of Growth Factor Receptors. In: Vande Woude GF, Klein G, eds. Vol 77. Advances in Cancer Research. Academic Press; 1999:25-79. 10.1016/S0065-230X(08)60784-8

11. Ménard S, Tagliabue E, Campiglio M, Pupa SM. Role of HER2 gene overexpression in breast carcinoma. J Cell Physiol. 2000;182(2):150–162. doi:10.1002/(SICI)1097-12.4652(200002)182:2<150::AID-JCP3>3.0.CO;2-E

12. HARRIS, Jay R. et al. Diseases of the Breast. 3rd ed. Ph. Lippincott Williams & Wilkins; 2012.

13. Burgess AW, Cho HS, Eigenbrot C, et al. An open-and-shut case? Recent insights into the activation of EGF/ErbB receptors. Mol Cell. 2003;12(3):541–552. doi:10.1016/S1097-2765(03)00350-2

14. Graus-Porta D, Beerli RR, Daly JM, Hynes NE. ErbB-2, the preferred heterodimerization partner of all ErbB receptors, is a mediator of lateral signaling. EMBO J. 1997;16(7):1647–1655. doi:10.1093/emboj/16.7.1647

15. Prenzel N, Fischer OM, Streit S, Hart S, Ullrich A. The epidermal growth factor receptor family as a central element for cellular signal transduction and diversification. Endocr Relat Cancer. 2001;8(1):11–31. doi:10.1677/erc.0.0080011

16. Ejskjær K, Sørensen BS, Poulsen SS, Forman A, Nexø E, Mogensen O. Expression of the epidermal growth factor system in endometrioid endometrial cancer. Gynecol Oncol. 2007;104(1):158–167. 10.1016/j.ygyno.2006.07.015

17. Edwards J, Traynor P, Munro AF, Pirret CF, Dunne B, Bartlett JMS. The role of HER1-HER4 and EGFRvIII in hormone-refractory prostate cancer. Clin Cancer Res. 2006;12(1):123–130. doi:10.1158/1078-0432.CCR-05-1445

18. Baker CH, Pino MS, Fidler IJ. Phosphorylated epidermal growth factor receptor on tumor-associated endothelial cells in human renal cell carcinoma is a primary target for therapy by tyrosine kinase inhibitors. Neoplasia. 2006;8(6):470–476. doi:10.1593/neo.06172

19. Ogiso H, Ishitani R, Nureki O, et al. Crystal structure of the complex of human epidermal growth factor and receptor extracellular domains. Cell. 2002;110(6):775–787. doi:10.1016/S0092-8674(02)00963-7

20. Hubbard SR. Structural analysis of receptor tyrosine kinases. Prog Biophys Mol Biol. 1999;71(3- 4):343–358. doi:10.1016/s0079-6107(98)00047-9

21. Keshamouni VG, Mattingly RR, Reddy KB. Mechanism of 17-β-Estradiol-induced Erk1/2 Activation in Breast Cancer Cells: A ROLE FOR HER2 AND PKC-δ*. J Biol Chem. 2002;277(25):22558–22565. 10.1074/jbc.M202351200

22. Rusnak DW, Affleck K, Cockerill SG, et al. The characterization of novel, dual ErbB-2/EGFR, tyrosine kinase inhibitors: potential therapy for cancer. Cancer Res. 2001;61(19):7196–7203.

23. Olayioye MA. Update on HER-2 as a target for cancer therapy: intracellular signaling pathways of ErbB2/HER-2 and family members. Breast Cancer Res. 2001;3(6):385–389. doi:10.1186/bcr327

24. Steven Grant, Liang Qiao PD. Roles of ERBB family receptor tyrosine kinases and downstream signaling pathways in the control of cell growth and survival. Pharmacology. 2002;(12):376–389.

25. Du Z, Lovly CM. Mechanisms of receptor tyrosine kinase activation in cancer. Mol Cancer. 2018;17(1):1–13. doi:10.1186/s12943-018-0782-4

26. Liu H, Zang C, Fenner MH, Possinger K, Elstner E. PPARgamma ligands and ATRA inhibit the invasion of human breast cancer cells in vitro. Breast Cancer Res Treat. 2003;79(1):63–74. doi:10.1023/a:1023366117157

27. Chavez KJ, Garimella S V, Lipkowitz S. Triple negative breast cancer cell lines: one tool in the search for better treatment of triple negative breast cancer. Breast Dis. 2010;32(1-2):35–48. doi:10.3233/BD-2010-0307

28. Lattrich C, Juhasz-Boess I, Ortmann O, Treeck O. Detection of an elevated HER2 expression in MCF-7 breast cancer cells overexpressing estrogen receptor β1. Oncol Rep. 2008;19(3):811–817. doi:10.3892/or.19.3.811

29. Hicks DG, Buscaglia B, Goda H, et al. A novel detection methodology for HER2 protein quantitation in formalin-fixed, paraffin embedded clinical samples using fluorescent nanoparticles: An analytical and clinical validation study. BMC Cancer. 2018;18(1):1–15. doi:10.1186/s12885-018-5172-1

30. Abramczyk H, Brozek-Pluska B, Kopec M, Surmacki J, Błaszczyk M, Radek M. Redox imbalance and biochemical changes in cancer by probing redox-sensitive mitochondrial cytochromes in label-free visible resonance raman imaging. Cancers (Basel). 2021;13(5). doi:10.1101/2020.12.03.409359

31. Abramczyk Halina, Beata Brozek-Pluska MK. Double face of cytochrome c in cancers by Raman imaging. Sci Rep. Published online 2022:1–11. doi:10.1038/s41598-022-04803-0

32. Dieing, T., Ibach W. Software Requirements and Data Analysis in Confocal Raman Microscopy. In Confocal Raman Microscopy. Published online 2011:61-89.

33. https://www.sigmaaldrich.com/life-science/proteomics/post-translational-analysis/phosphorylation/protein-phosphorylation-analysis-tools.html. https://covid19.who.int/

34. Kalpage HA, Wan J, Morse PT, et al. Cytochrome c phosphorylation: Control of mitochondrial electron transport chain flux and apoptosis. Int J Biochem Cell Biol. 2020;121:105704. 10.1016/j.biocel.2020.105704

35. https://www.thermofisher.com/pl/en/home/life-science/protein-biology/protein-biology-learning-center/protein-biology-resource-library/pierce-protein-methods/phosphorylation.html.

36. Okishio N, Fukuda R, Nagai M, Nagai Y, Nagatomo S, Kitagawa T. Tyrosine Phosphorylation-induced Changes in Absorption and UV Resonance Raman Spectra of Src-Peptides. J Raman Spectrosc. 1998;29(1):31–39. doi:10.1002/(sici)1097-4555(199801)29:<31::aid-jrs208>3.0.co;2-b

37. Stone N, Kendall C, Smith J, Crow P, Barr H. Raman spectroscopy for identification of epithelial cancers. Faraday Discuss. 2004;126(1):141–157. doi:10.1039/b304992b

38. Thomas GJ. Raman spectroscopy of protein and nucleic acid assemblies. Annu Rev Biophys Biomol Struct. 1999;28:1–27. doi:10.1146/annurev.biophys.28.1.1

39. Shaver JM, Christensen KA, Pezzuti JA, Morris MD. Structure of Dihydrogen Phosphate Ion Aggregates by Raman-Monitored Serial Dilution. Appl Spectrosc. 1998;52(2):259–264. https://opg.optica.org/as/abstract.cfm?URI=as-52-2-259

40. Xie Y, Zhang D, Jarori GK, Davisson VJ, Ben-Amotz D. The Raman detection of peptide tyrosine phosphorylation. Anal Biochem. 2004;332(1):116–121. doi:10.1016/j.ab.2004.05.052

41. Deng H, Bloomfield VA, Benevides JM, Thomas GJ. Dependence of the raman signature of genomic B-DNA on nucleotide base sequence. Biopolymers. 1999;50(6):656–666. doi:10.1002/(SICI)1097-0282(199911)50:6<656::AID-BIP10>3.0.CO;2-9

42. Guan Y, Thomas GJ. Vibrational analysis of nucleic acids. IV. Normal modes of the DNA phosphodiester structure modeled by diethyl phosphate. Biopolymers. 1996;39(6):813–835. doi:10.1002/(sici)1097-0282(199612)39:6<813::aid-bip7>3.0.co;2-y

43. Guan Y, Choy GSC, Glaser R, Thomas GJ. Vibrational analysis of nucleic acids. 2. Ab initio calculation of the molecular force field and normal modes of dimethyl phosphate. J Phys Chem. 1995;99(31):12054-12062. doi:10.1021/j100031a039

44. Wilkinson GRRJHC and REH. Advances in Infrared and Raman Spectroscopy. Published online 1985:487.

45. Abramczyk H, Surmacki J, Kopeć M, Olejnik AK, Lubecka-Pietruszewska K, Fabianowska-Majewska K. The role of lipid droplets and adipocytes in cancer. Raman imaging of cell cultures: MCF10A, MCF7, and MDA-MB-231 compared to adipocytes in cancerous human breast tissue. Analyst. 2015;140(7):2224–2235. doi:10.1039/c4an01875c

46. Spiro TG, Strekas TC. Resonance Raman Spectra of Heme Proteins. Effects of Oxidation and Spin State. J Am Chem Soc. 1974;96(2):338–345. doi:10.1021/ja00809a004

47. Hu S, Spiro TG, Morris IK, Singh JP, Smith KM. Complete Assignment of Cytochromec Resonance Raman Spectra via Enzymatic Reconstitution with Isotopically Labeled Hemes. J Am Chem Soc. 1993;115(26):12446–12458. doi:10.1021/ja00079a028

48. Wood BR, McNaughton D. Raman excitation wavelength investigation of single red blood cells in vivo. J Raman Spectrosc. 2002;33(7):517–523. doi:10.1002/jrs.870

49. Abe M, Kitagawa T, Kyogoku Y. Resonance Raman spectra of octaethylporphyrinato-Ni(II) and meso-deuterated and 15N substituted derivatives. II. A normal coordinate analysis. J Chem Phys. 1978;69(10):4526–4534. doi:10.1063/1.436450

50. Hu S, Smith KM, Spiro TG. Assignment of protoheme Resonance Raman spectrum by heme labeling in myoglobin. J Am Chem Soc. 1996;118(50):12638–12646. doi:10.1021/ja962239e

51. Abramczyk H, Surmacki JM, Brozek-Pluska B. Redox state changes of mitochondrial cytochromes in brain and breast cancers by Raman spectroscopy and imaging. J Mol Struct. 2022;1252:132134. 10.1016/j.molstruc.2021.132134

52. Tai W, Mahato R, Cheng K. The role of HER2 in cancer therapy and targeted drug delivery. J Control Release. 2010;146(3):264–275. 10.1016/j.jconrel.2010.04.009

53. Tanos T, Rojo LJ, Echeverria P, Brisken C. ER and PR signaling nodes during mammary gland development. Breast Cancer Res. 2012;14(4). doi:10.1186/bcr3166

54. Perou CM, Sørlie T, Eisen MB, et al. Molecular portraits of human breast tumours. Nature. 2000;406(6797):747-752. doi:10.1038/35021093

55. Sørlie T, Perou CM, Tibshirani R, et al. Gene expression patterns of breast carcinomas distinguish tumor subclasses with clinical implications. Proc Natl Acad Sci U S A. 2001;98(19):10869–10874. doi:10.1073/pnas.191367098

56. Sørlie T, Tibshirani R, Parker J, et al. Repeated observation of breast tumor subtypes in independent gene expression data sets. Proc Natl Acad Sci U S A. 2003;100(14):8418–8423. doi:10.1073/pnas.0932692100

57. Subik K, Lee J-F, Baxter L, et al. The Expression Patterns of ER, PR, HE R2, CK5/6, EGFR, Ki-67 and AR by Immunohistochemical Analysis in Breast Cancer Cell Lines. Breast Cancer (Auckl). 2010;4:35-41.

58. Abramczyk H, Surmacki JM, Kopeć M, Jarczewska K, Romanowska-Pietrasiak B. Hemoglobin and cytochrome c. reinterpreting the origins of oxygenation and oxidation in erythrocytes and in vivo cancer lung cells. Sci Rep. 2023;13(1):1–15. doi:10.1038/s41598-023-41858-z

